# Increased frequency of travel in the presence of cross-immunity may act to decrease the chance of a global pandemic

**DOI:** 10.1101/404871

**Authors:** R.N. Thompson, C.P. Thompson, O. Pelerman, S. Gupta, U. Obolski

**Affiliations:** Mathematical Institute, University of Oxford, Andrew Wiles Building, Radcliffe Observatory Quarter, Woodstock Road, Oxford OX2 6GG, UK; Department of Zoology, University of Oxford, South Parks Road, Oxford OX1 3PS, UK; Christ Church, University of Oxford, St Aldate’s, Oxford OX1 1DP, UK; The Chaim Rosenberg School of Jewish Studies, Tel Aviv University, Tel Aviv 69978, Israel

**Keywords:** major epidemic, antigenic variation, cross-immunity, pathogen diversity, mathematical modelling

## Abstract

The high frequency of modern travel has led to concerns about a devastating pandemic since a lethal pathogen strain could spread worldwide quickly. Many historical pandemics have arisen following pathogen evolution to a more virulent form. However, some pathogen strains invoke immune responses that provide partial cross-immunity against infection with related strains. Here, we consider a mathematical model of successive outbreaks of two strains – a low virulence strain outbreak followed by a high virulence strain outbreak. Under these circumstances, we investigate the impacts of varying travel rates and cross-immunity on the probability that a major epidemic of the high virulence strain occurs, and the size of that outbreak. Frequent travel between subpopulations can lead to widespread immunity to the high virulence strain, driven by exposure to the low virulence strain. As a result, major epidemics of the high virulence strain are less likely, and can potentially be smaller, with more connected subpopulations. Cross-immunity may be a factor contributing to the absence of a global pandemic as severe as the 1918 influenza pandemic in the century since.

## 1. INTRODUCTION

Outbreaks of infectious disease are responsible for around 14 million deaths annually [1,2]. In recent years, there have been a number of epidemics that have sparked fears that a global pandemic might develop [3]. Outbreaks such as the 2013-16 Ebola epidemic in West Africa did not develop into a global pandemic [4]. Other outbreaks, for example the 2009 H1N1 influenza pandemic, have included cases in countries worldwide [5] but have not been as destructive as initially feared [6]. Historically, however, there have been severe global pandemics. The 1918 ‘Spanish flu’ pandemic killed around 50 million people [7], and the 1958 and 1968 influenza pandemics caused around one million deaths each [8,9]. These outcomes viewed collectively raise a number of questions: how likely is a global pandemic now, and has the lack of a devastating global pandemic in recent years been simply a matter of luck?

A number of factors affect the pandemic potential when a pathogen first appears in a host population. These include the spatial distribution of hosts and the level of mixing between subpopulations [10,11]. When a pathogen enters a population, a high host density within subpopulations and a high contact rate between subpopulations are most likely to represent appropriate conditions for a major epidemic and lead to high epidemic growth rates [12-15]. The modern world increasingly satisfies these conditions, with growing population sizes and large numbers of individuals living in urban centres, as well as the high frequency of worldwide airline travel [16]. Hence, one might expect the probability of a global pandemic, as well as its potential severity, to be at an all-time high.

However, the determinants of the dynamics of infectious disease outbreaks are numerous and the above assertions ignore an important feature: cross-immunity obtained from pathogen exposures in previous outbreaks. Viral, bacterial and eukaryotic pathogens, such as the influenza virus, *Streptococcus pneumoniae* and the malaria parasite, evolve in an attempt to avoid control by the immune system [17,18] and in response to interventions such as vaccines [19-21]. Nonetheless, such immune evasion is not perfect. Hosts that have been infected by a particular strain and acquired immunity to that strain may also be at least partially immune to infection with related strains [22-26]. Previous infections with immunologically related strains of a pathogen can therefore be beneficial to hosts as they might provide protection against future infections with other potentially more virulent strains.

Cross-immunity is known to affect the dynamics of outbreaks of various pathogens. *Vibrio cholera*, for example, exhibits almost complete cross-immunity between strains (a study by Koelle *et al.* [27] used an estimate that prior infection by a related strain reduces the infection rate by 95.6%). Human Respiratory Syncytial Virus (hRSV) has been shown to provide incomplete cross-immunity against infections from the same virus group [28], and low levels of cross-immunity again hRSV infections from different virus groups [28]. Furthermore, it provides partial cross-immunity against infections with human Parainfluenza Virus [29].

Cross-immunity between related pathogens or pathogen strains has also been shown to impact on pathogen dynamics and the structures of pathogen populations [30,31]. Cross-immunity might also be expected to affect the chance of and threat from a major epidemic. The 2009 H1N1 pandemic, for example, was not as destructive as feared, potentially as a result of decreased population susceptibility due to cross-immunity [32-35]. Here we develop a mathematical model to investigate the impact of cross-immunity on the chance of a major epidemic. We consider a general system emulating the dynamics of an outbreak of a pathogen of high virulence in two connected subpopulations when cross-immunity is present from a previously circulating low virulence strain. We illustrate the principle that high rates of travel between subpopulations can decrease the probability of a major epidemic of the high virulence strain, and that the expected outbreak size can be either increased or decreased when there is a higher rate of travel between subpopulations. In a connected world, novel pathogens could spread worldwide through immunologically naïve populations extremely quickly. However, epidemics of existing pathogens may be less frequent and potentially smaller due to cross-immunity between pathogen strains – an important, but previously underappreciated, factor.

## 2. RESULTS

Motivated by the spread of pathogens between geographically separate regions, we considered pathogen transmission in a population consisting of two spatially distinct subpopulations (Fig 1). Two outbreaks were assumed to occur. First, an outbreak of a low virulence (LV) strain of the pathogen that has the potential to generate a large number of cases but that is unlikely to lead to a large number of deaths. Then, an outbreak of a related high virulence (HV) strain of the pathogen. Individuals infected in the first outbreak were partially immune against infection in the second outbreak. It is the probability of a major epidemic of the HV strain and the possible final sizes of the HV strain outbreak that are our main concerns in this article.

**Figure 1.**
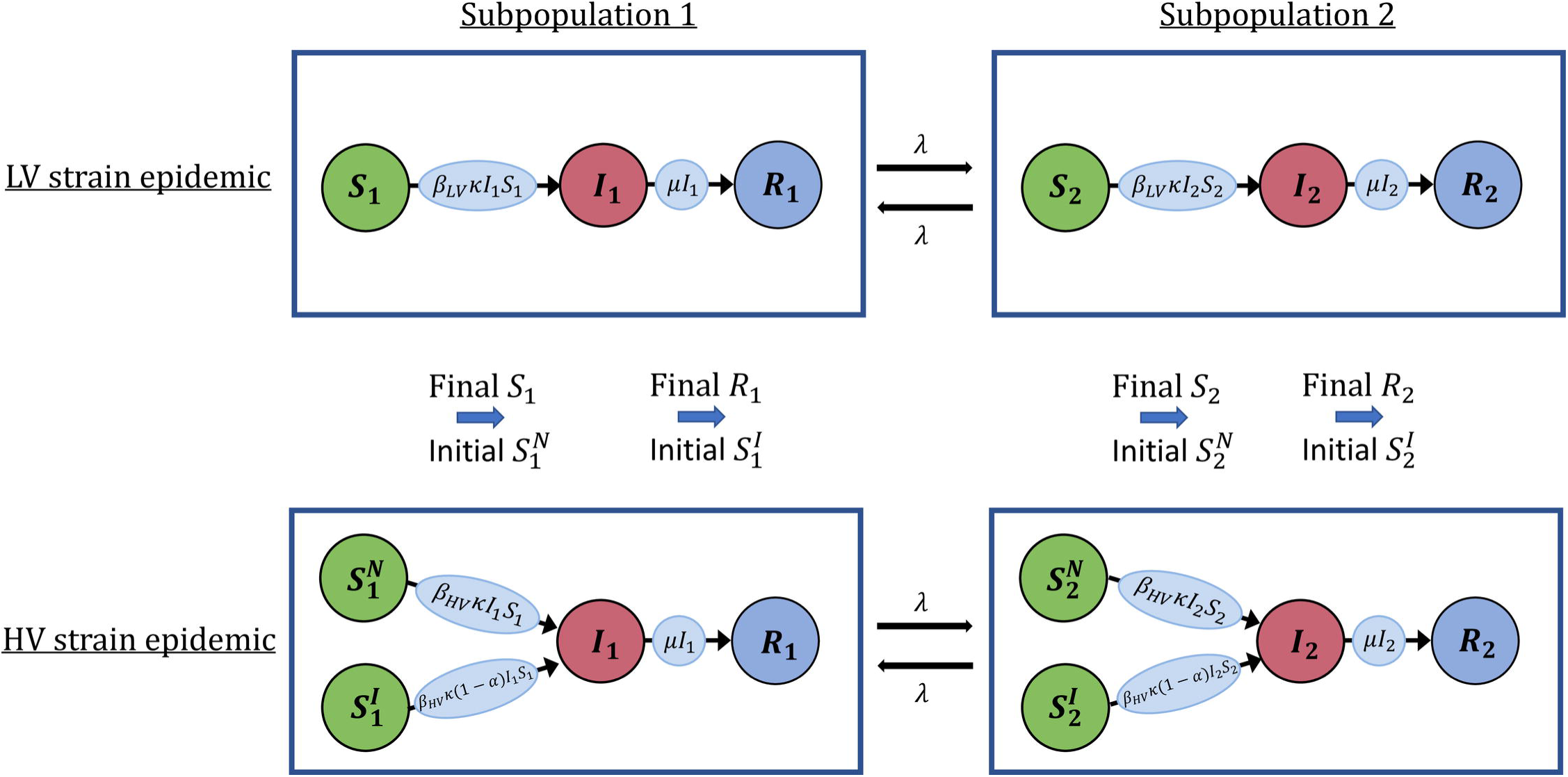
Schematic illustrating how outbreaks are simulated using the two-subpopulation model. Circles represent susceptible (*S*_1_, *S*_2_), infected (*I*_1_, *I*_2_) and removed (*R*_1_, *R*_2_) individuals in subpopulations 1 and 2, while the rates of transfer of individuals between these classes are given on the arrows connecting the circles (see Methods and Supplementary Material). A major epidemic of the LV strain is assumed to occur, followed by an outbreak of the HV strain. Individuals infected in the LV strain epidemic are conferred cross-immunity at a level α against infection with the HV strain.

### Outbreak dynamics in a single subpopulation

We initially considered outbreak dynamics in a single population. When the HV strain arrived in the population, its effective reproduction number was reduced if individuals had previously been infected by the LV strain. The higher the cross-immunity (governed by the level of cross-immunity, 0*≤ α ≤* 1), the lower the effective reproduction number of the HV strain (Fig 2a). The higher the basic reproduction number of the LV strain, 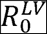, the lower the expected effective reproduction number of the HV strain. This is because, when the LV strain had a high basic reproduction number, it generated a large number of cases, thereby leading to large numbers of individuals with partial cross-immunity against the HV strain (Fig 2a).

**Figure 2.**
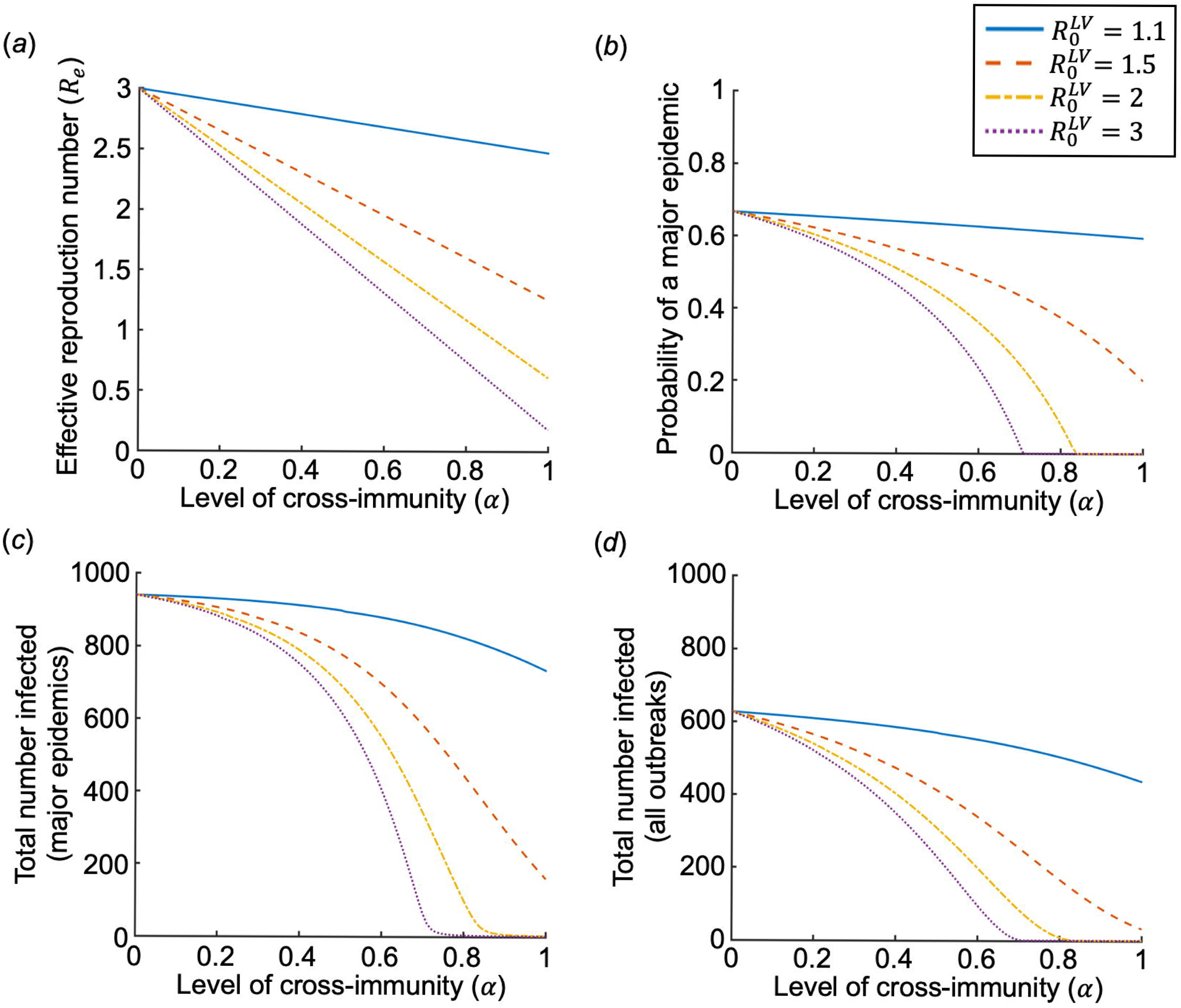
The impact of cross-immunity on outbreaks of the HV strain in a single population. (a) The effective reproduction number of the HV strain (α_e_); (b) The probability of a major epidemic of the HV strain; (c) The final size of the HV strain outbreak when a major epidemic occurs 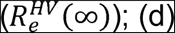 The expected total number of individuals infected by the HV strain, allowing for the possibility of a major epidemic or fade-out without causing a major epidemic. The level of cross-immunity (α) is displayed on the x-axis, and lines represent different values of the basic reproduction number of the LV strain, 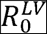. A major epidemic of the LV strain was assumed to occur prior to the arrival of the HV strain in the population. Outbreaks of the HV strain were seeded with a single infected individual, with all other individuals susceptible (either immunologically naïve, 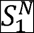, or cross-immune following infection with the LV strain, 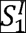). Parameter values: *N* = 1000, κ= 1, 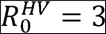. For a description of the parameters, see Table S2.

Similarly, when the HV strain arrived in the population, the probability of a major epidemic (i.e. successful invasion of the host population) decreased as the level of cross-immunity increased (Fig 2b), as did the final size of major epidemics of the HV strain (Fig 2c). The expected number of HV strain infections, accounting for the possibility that outbreaks may fade-out without becoming major epidemics, also decreased when the level of cross-immunity increased (Fig 2d). Furthermore, these quantities were also all reduced when the LV strain was more transmissible (see different lines in Figs 2b-d).

We also considered stochastic simulations of the model (Fig 3), which support the analytic result that cross-immunity lowers the probability of a major epidemic of the HV strain in addition to its final size. The results in Figs 3b-c can be compared with Figs 2b-c: for *α* = 0 and *α* = 0.5, the probability of a major epidemic in Fig 2b when 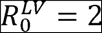 = 2 takes the values 0.68 and 0.43, respectively – matching the percentage of outbreaks that are major epidemics in Figs 3b and 3c. Similarly, for *α* = 0 and *α* = 0.5, the total numbers of infections in Fig 2c when 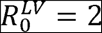 are 940 and 693, respectively – which are also consistent with the major epidemics simulated in Figs 3b-c.

**Figure 3.**
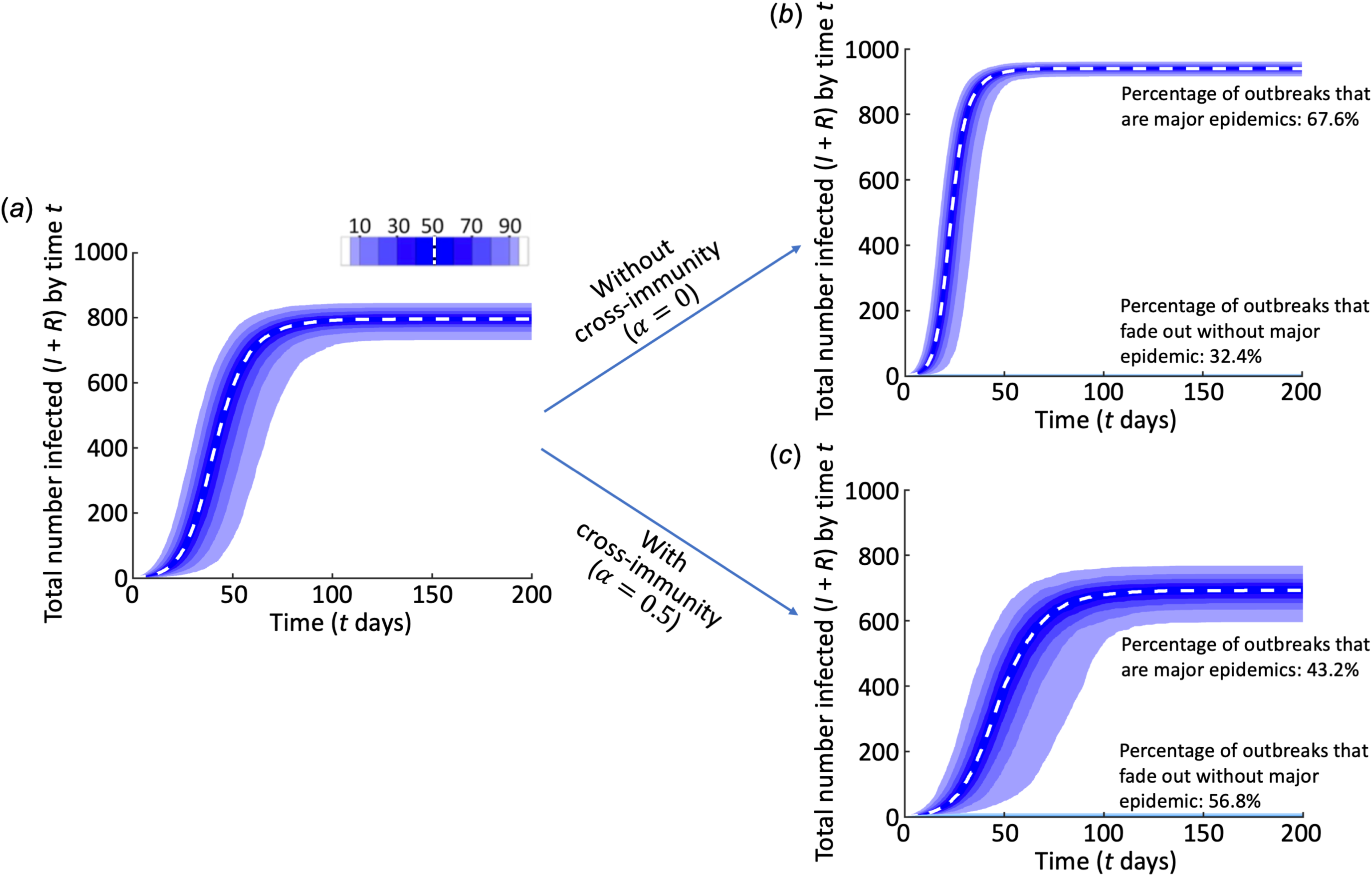
Cross-immunity leads to fewer and smaller major epidemics. Simulations of the model in a single population showing: (a) The LV strain major epidemic; (b) The subsequent HV strain outbreak, if there is no cross-immunity (α 0); (c) As in panel b, but with cross-immunity (α 0.5). Parameter values: *N* 1000, κ = 1,µ 0.143 day^−1^β_LV_ 2.86×10^−4^ day^−1^ (so that 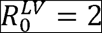), β_HV_ 4.29×10^−4^ day^−1^ (so that 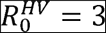). For a description of the parameters, see Table S2. The blue regions represent quantiles of simulations of major epidemics and the white dotted line represents the median values. In panels b-c, there are also thin light blue areas at the bottoms of the panels showing outbreaks that fade-out without causing major epidemics. The light blue area is omitted in panel a, since we only consider major epidemics of the LV strain. Simulations are run in the same fashion as in the case of two connected subpopulations but with the rate of travel between subpopulations set to λ= 0, and with the outbreaks always seeded with a single infected individual in the first subpopulation with the remaining *N –* 1 individuals susceptible. In the HV strain epidemic, each susceptible individual is either immunologically naïve and in the class 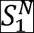, or cross-immune following infection with the LV strain and in the class 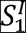.

### Outbreak dynamics in the two-subpopulation model

We then considered outbreak dynamics in two connected subpopulations and assessed the impact of the extent of travel between subpopulations (governed by the parameter *λ*), as well as the within-subpopulation transmissibility of the LV and HV strains (governed by the parameter *κ*), on outbreaks of the HV strain. We considered the effects of these parameters on the probability of a major epidemic of the HV strain (Fig 4a-c), the final size of an outbreak if a major epidemic is assumed to occur (Fig 4d-f), and the expected final size accounting for the fact that outbreaks can either fade out as minor outbreaks or take off as major epidemics (Fig 4g-i). We found that increased rates of travel between subpopulations led to a lower probability of a major epidemic of the HV strain. This is due to the probability of a major epidemic of the HV strain being governed by local susceptibility when the HV strain first arrives in the population. Thus, increased travel between subpopulations increases the probability that the LV strain has previously caused a major epidemic in the subpopulation where the HV strain arrives and conferred some immunity against the HV strain in that subpopulation. The decreased chance of a major epidemic of the HV strain when between-subpopulation travel was increased was particularly pronounced when the strains driving the LV and HV epidemics were very transmissible and the level of cross-immunity was high (see rightmost region of Fig 4c, in which there is a large reduction in the probability of a major epidemic of the HV strain from the bottom to the top of the figure).

**Figure 4.**
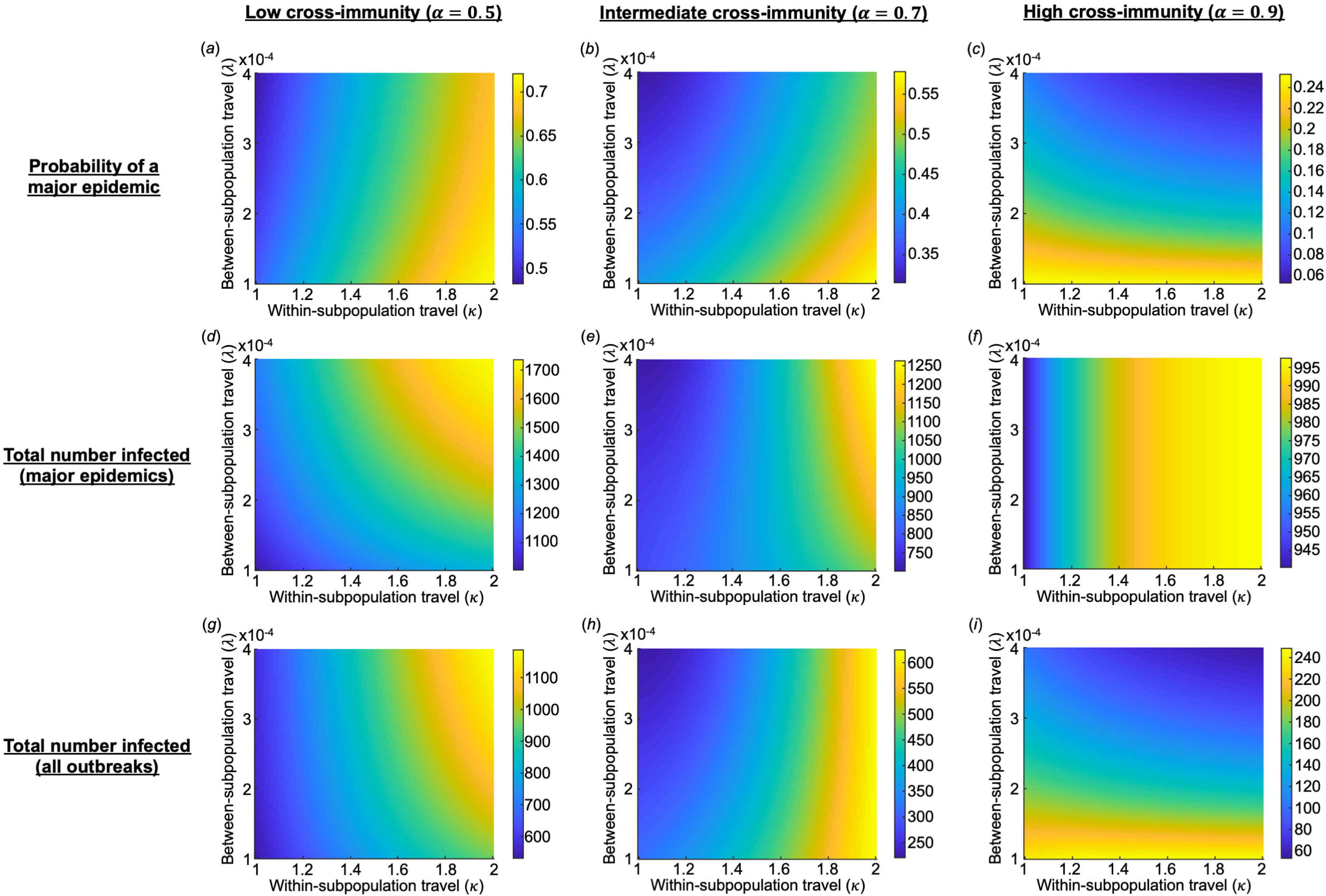
The impact of within-and between-subpopulation travel on outbreaks of the HV strain in the two-subpopulation model. The left, middle and right columns are results with the level of cross-immunity, *α*, equal to 0.5, 0.7 and 0.9, respectively. (a)-(c): the probability of a major epidemic of the HV strain; (d)-(f): the expected final size of major epidemics of the HV strain; (g)-(i): the expected total number of the HV strain infections, allowing for the possibility of a major epidemic or fade-out without causing a major epidemic. A major epidemic of the LV strain is assumed to occur in at least one subpopulation and is seeded by a single infected individual in subpopulation 1 (see Fig 1 and Supplementary Material). Outbreaks of the HV strain are seeded with a single infected individual in a subpopulation chosen at random, with all other individuals in both subpopulations susceptible. Parameter values: N = 1000 individuals per subpopulation, within-subpopulation basic reproduction numbers 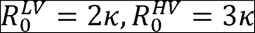. For a description of the parameters, see Table S2.

If the between-subpopulation connectivity was high when major epidemics of the HV strain occurred, those epidemics tended to be more severe than when the between-subpopulation connectivity was lower (Fig 4d-e). This was not the case at very high levels of cross-immunity (Fig 4f). However, the expected total number of HV strain infections, accounting for the possibility that the outbreak either took off and became a major epidemic or faded out without spreading widely, reduced as the extent of between-subpopulation travel increased when the level of cross-immunity was high (e.g. Fig 4h-i) but increased for lower levels of cross-immunity (Fig 4g).

The outcome that the expected final size of major epidemics of the HV strain was unaffected by between-subpopulation travel when the level of cross-immunity was high (Fig 4f) can be explained as follows. The LV strain was assumed to always generate a major epidemic in subpopulation 1. In that subpopulation, the cross-immunity conferred was so high that a major epidemic of the HV strain could never occur in that subpopulation. Consequently, the only way that a major epidemic of the HV could occur in the overall population was if a major epidemic of the LV strain did not occur in subpopulation 2, and the HV strain outbreak was seeded in subpopulation 2 and went on to cause a major epidemic. Conditional on this scenario occurring, the size of that major epidemic would be unaffected by the rate of between-subpopulation travel, as can be seen in Fig 4f.

We also verified that our results in Fig 4, which were derived numerically for computational efficiency and based on analytical expressions (see Supplementary Material), matched the results of stochastic simulations (Fig S1).

## 3. DISCUSSION

The large increase in international travel over the last century might be assumed to have resulted in a high chance of a devastating global pandemic (see e.g. [36]). Here we have used a general epidemiological model to demonstrate that an important, yet often overlooked, factor in the dynamics of a newly introduced high-virulence (HV) pathogen strain is partial immunity driven by exposures to related pathogen strains. When a HV pathogen strain arrives in a population following an epidemic of a related but low virulence (LV) strain, the probability of a major epidemic of the HV strain is decreased. High rates of travel between spatially distinct subpopulations can drive larger outbreaks of low virulence pathogens, in turn providing higher levels of immunity if/when a HV strain, which has the potential to cause a devastating epidemic, appears in the population (Fig 4a-c).

Not only did we find that the probability of a major epidemic of the HV strain decreases when travel between subpopulations increases but the expected final size of the HV strain outbreak can also be reduced. This was particularly pronounced when the level of cross-immunity between strains was high (Fig 4i) since lower cross-immunity levels combined with high travel rates can lead to large epidemics due to increased mixing between subpopulations (Fig 4g). When between-subpopulation travel was increased, the reduction in the probability of a major epidemic of the HV strain, and the reduction in the expected size of the HV strain outbreak, was largely due to cross-immunity reducing the proportion of outbreaks that proceeded to become major epidemics. When major epidemics occurred, we found that they were typically larger when there was more travel between regions (Fig 4d-e), although this was not always the case, particularly when the level of cross-immunity was very high (Fig 4f).

Partial cross-immunity against a highly virulent strain from prior exposure to a less virulent strain is characteristic of influenza epidemics. For example, it has been suggested that individuals born before 1890 were protected against the 1918 H1N1 pandemic due to the outbreak in 1889-90 [37] and that individuals infected with multiple historical seasonal H1N1 influenza strains were protected against the 2009 H1N1 influenza pandemic strain [38]. Potentially related to this, evidence indicates that older adults are less likely to be severely affected by influenza pandemics than younger adults, and this could be because of cross-immunity from exposure to related strains [39]: in the 1977-78 H1N1 epidemic, for example, only adults under 26 years of age experienced substantial morbidity [40,41]. Not only does cross-immunity protect individuals from infection, but our results also suggest that cross-immunity might be a potential explanatory factor as to why there has not been a pandemic as devastating as the 1918 influenza epidemic in the century since, despite the emergence of a strain antigenically similar to the 1918 pandemic strain in 2009. Increased travel during this time period may have led to more widespread cross-immunity against strains of pandemic potential, due to the global spread of related, less virulent strains. To verify that travel rates have indeed increased substantially during the 20^th^ century, as a theoretical exercise we obtained crude estimates of the travel rates from Europe to the USA, comparing rates during the early 20th century with current travel rates. For the early 20^th^ century rates, we used registry statistics from USA ports from 1914 to 1924 (see Supplementary Material Section 5). Oceanic travel should approximate overall trans-Atlantic travel in this time period, as other modes of long-distance travel were rare. This yielded the approximation *λ* ≈ 3.6· 10^−6^ per day. Conversely, when estimating current rates of travel from European countries to the USA, approximated using data on air travel, we obtain much higher values of *λ* ≈ 3.7· 10^−4^ per day. It might be expected that such a significant rise in travel in this time period might have reduced the risk of a global pandemic of a pathogen with circulating strains that induce cross-immunity (Fig 4).

Our assertion that higher rates of travel in the presence of cross-immunity may reduce the impact of an epidemic is supported by evidence that island nations are more affected during influenza pandemics than those that are connected by land to other countries. Isolated islands such as the Solomon Islands experienced mortality rates of in the range 4-10% during the 1918 pandemic [42], whereas more connected regions of the world that had experienced more recent epidemics saw a mortality rate of 1-3% [43]. Furthermore, with respect to morbidity, we found that out of the 30 countries with the highest number of cases per capita during the 2009 H1N1 pandemic, 24 were island nations [44].

While we have chosen to focus on pathogens affecting human populations, we note that cross-immunity might also impact on the pandemic potential for pathogens of animals and plants. Cross-immunity between pathogen strains is a feature of pathogens of animals including *Trypanosoma congolense* [45] and pathogens of plants including Tobacco mosaic virus and Potato virus X [46]. Moreover, a previous study of the plant pathogen *Podosphaera plantaginis* found that highly connected populations experienced less pathogen colonisation and higher pathogen extinction than populations that were less connected, and attributed this to the evolution of disease resistance [47]. Furthermore, the effect of cross-immunity on plant or animal disease outbreaks might also be changing due to alterations in the worldwide movement of hosts. For example, climate change appears to be modifying the spatial distributions of animal populations [48] and the plant nursery trade is more active than ever before [49]. We also note that, while we focussed on cross-immunity between different pathogen strains, a related concept is protection against reinfection with the same (or very similar) strain – often referred to as homologous interference or superinfection exclusion – which has been demonstrated for a number of pathogens of plants or animals [50].

We aimed here to develop the simplest model possible characterising the spread of a pathogen between spatially distinct populations in the presence of cross-immunity. For assessing the probability of a major epidemic of a specific virulent strain of a pathogen in a particular host population, the model would need to be extended and adjusted. The process of mutation of the pathogen from the low virulence strain to the high virulence strain may have to be modelled explicitly. If the mutation occurs a long period after the previous major epidemic of the low virulence strain, cross-immunity might be expected to have waned compared to if the low and high virulence strain epidemics occur in quick succession. Alternatively, the high virulence strain epidemic might start before the low virulence strain epidemic has ended. This would require an extension to the epidemiological model, to permit the low and high virulence strains to co-circulate (for a preliminary analysis, see Supplementary Information Section 6). The cumulative effects of a number of past outbreaks of different related strains might also have to be considered when a high virulence strain enters a host population [25,38]. Different types of partial cross-immunity could be considered. Here we have assumed that exposure to the LV strain reduces the probability of infection with the HV strain, whereas for some infections cross-immunity may instead (or also) act to reduce the severity of disease [41,51,52]. As an example, epidemiological isolation leading to high mortality rates of infections may explain the mass mortality in pacific island populations between the 16^th^ and 19^th^ centuries [53].

Other extensions to the model might have to be considered for application to specific epidemics in human, animal or plant populations. For example, to explore the contrasting effects of cross-immunity (which protects previously infected hosts) and antibody-dependent enhancement (which would be expected to increase disease severity in previously infected hosts [54]) for dengue epidemics in human populations, a detailed model conditioned to the epidemiology of dengue would have to be developed. In doing this, however, we note that incorporation of additional epidemiological detail in models is not always useful, and that only necessary features should be included to allow model outputs to be clearly understood [55]. For all epidemics, control interventions might have to be included in the model, such as movement bans [56] or trade restrictions [57] for livestock or commercial plant diseases. These particular interventions could be included in the model by altering the within-subpopulation or between-subpopulation rates of travel. Linked to this, behavioural responses of individuals in the host population might have to be considered even if interventions are not in place – for example, the awareness of the risk posed by the pathogen might be higher following the LV strain epidemic, leading to reduced travel during the HV strain epidemic.

Nonetheless, we have demonstrated the principle that increased global travel might not necessarily mean that large pandemics are more likely in the present day than previously. On the contrary, our results demonstrate that there may exist conditions under which increased travel between subpopulations might reduce the probability and size of major epidemics. We do not contest that an increase in travel may sometimes increase the chance of a major epidemic, as well as its final size. The size of an epidemic of an entirely new pathogen, for example, or a strain that is antigenically distinct from previously circulating strains, is likely to be larger when there is more travel. This is because the pathogen would then be entering a population that is immunologically naïve and additional mixing provides the opportunity for more transmission events. There is therefore a need to test whether or not increased travel leads to smaller or larger epidemics in different theoretical contexts, as well as using data from a range of real-world outbreaks. However, we have focussed on epidemics occurring due to variants of pre-existing pathogens since these have driven a substantial number of past pandemics. Predicting which pathogen is likely to cause the next major pandemic is challenging [58]. Since our results demonstrate how previous outbreaks may have spread cross-immunity against strains related to pre-existing pathogens, our study has led us to support the assertion of the World Health Organisation: the pandemic threat may be greatest from an unknown strain of a known pathogen, or a pathogen that we have not previously encountered, the so-called disease X [59].

## 4. METHODS

### Mathematical model

#### LV strain outbreak

We first considered an outbreak of a low virulence (LV) strain. Individuals were classified according to whether they were (*S*)usceptible to the LV strain, (*I*)nfected, or (*R*)ecovered and immune to that strain. Within a single subpopulation, the deterministic SIR model is given by

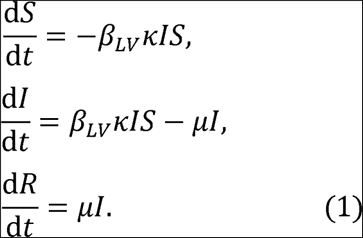

The parameter *κ* is a proxy for the frequency of within-subpopulation travel and governs the magnitude of the contact rate within the subpopulation. This parameter is also used in the subsequent high virulence (HV) strain outbreak (see below). The LV strain epidemic was assumed to follow stochastic SIR dynamics, analogous to the system of equations (1) within each subpopulation, with individuals also travelling between subpopulations at rate *λ* per day (see Fig 1 and Supplementary Material).

#### HV strain outbreak

We then considered a subsequent outbreak of the HV strain in the population, that is also governed by stochastic SIR dynamics in each subpopulation but extended to account for cross-immunity.

Individuals infected in the LV strain epidemic were partially protected against the HV strain epidemic. The extent of immunity against the HV strain conferred by the LV strain was governed by a parameter 0 ≤ *α* ≤ 1, which changed the probability of successful infection by a multiplicative factor 1-*λ* if the host had previously been infected by the LV strain compared to if the host had not previously been infected by the LV strain. The value *λ* 0 thereby corresponded to no cross-immunity, *λ* 1 corresponded to complete cross-immunity, and intermediate values of *λ* corresponded to partial cross-immunity.

Individuals were classified according to whether they were (*S*_N_)usceptible and not partially cross-immune against the HV strain, (*S*_1_)usceptible and partially cross-immune, (*I*)nfected, or (*R*)ecovered and immune to that strain or dead. The deterministic analogue in a single subpopulation to the model that we considered to represent the HV strain outbreak is

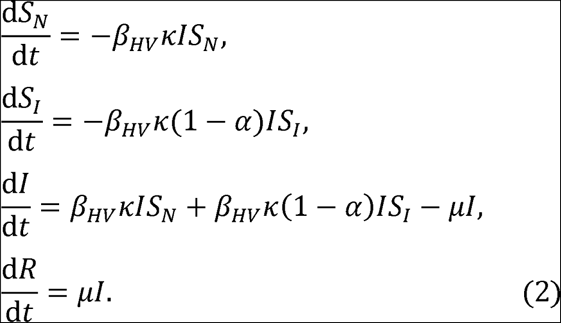

When the HV strain arrived in the subpopulation, initial cases could either lead to a major epidemic or fade-out as a minor outbreak. The within-subpopulation basic reproduction numbers of the LV and HV strains were denoted by 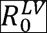 and 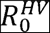, respectively (see Supplementary Material). In the HV strain outbreak, like in the previous LV strain outbreak, we also assumed that individuals travelled between subpopulations at rate *λ* per day (see Fig 1).

### Background theory

Before considering the full model, we derived the probability of a major epidemic and the expected final size conditional on a major epidemic when the pathogen first arrived in a subpopulation, ignoring any effects of migration between subpopulations.

#### LV strain epidemic

When the LV strain, with within-subpopulation basic reproduction number 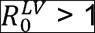, arrived in a single fully susceptible population, the probability of a major epidemic following was approximately

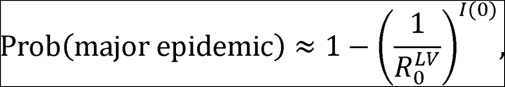

where *I*(0) is the number of infected hosts that were infected initially (see e.g. [60,61]).

If a major epidemic occurred, the expected number of individuals infected over the course of the epidemic was approximated by the solution, *R*^*LV*^(∞), of the final size equation (see equation (S3) in the Supplementary Material),

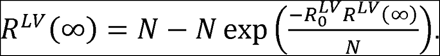

#### HV strain epidemic

When the HV strain appeared in a subpopulation, if a major epidemic of the LV strain had not occurred in that subpopulation, then the probability of a major epidemic was

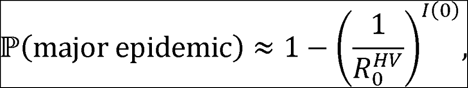

whenever 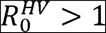, where *I*(0) is the initial number of hosts infected by the HV strain at the start of the HV strain outbreak. The final size if a major epidemic occurred was then given by the solution, 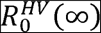, of the final size equation

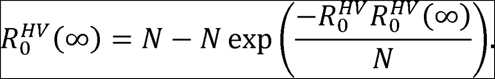

However, when the HV strain arrived in the subpopulation, if a major epidemic of the LV strain had occurred previously, then the population was not fully susceptible due to partial cross-immunity. The effective reproduction number of the HV strain when it arrived in the population was then 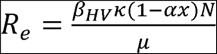, where 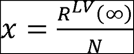 is the proportion of individuals infected in the LV strain epidemic (see Supplementary Material Section 2).

The resulting probability of a major epidemic of the HV strain was then

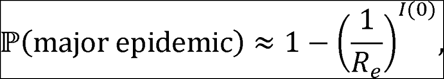

whenever *R*_e_ > 1, and if a major epidemic occurred, the final size was approximated by numerically solving the system of equations (2).

### Possible outcomes of LV and HV strain outbreaks in two subpopulations

When we considered the full model, consisting of two subpopulations (denoted 1 and 2), we assumed without loss of generality that the LV strain arrived in subpopulation 1. Since we were only interested in the effect of cross-immunity on outbreaks of the HV strain, we assumed that the LV strain successfully invaded the population. As a result, a major epidemic of the LV strain occurred in subpopulation 1.

Eight outcomes were then possible (see Table S1). A major epidemic of the LV strain may or may not occur in subpopulation 2. Then, major epidemics of the HV strain may or may not occur in each of subpopulations 1 and 2. Calculation of the probabilities of each of these outcomes, as well as the expected numbers of individuals infected by the LV and HV strains in each subpopulation, are described in the Supplementary Material.

## DATA AVAILABILITY

All numerical calculations and stochastic simulations in this manuscript were performed in Matlab. Accompanying code for the full model can be found at www.github.com/robin-thompson/PandemicPotentialGivenCrossProtection.

## AUTHORS’ CONTRIBUTIONS

RNT and SG conceived the research; All authors designed the study; RNT and UO carried out the research; RNT drafted the manuscript; All authors revised the manuscript and gave final approval for publication.

### ACKNOWLEDGEMENTS

Thanks to members of the Department of Zoology and Mathematical Institute in Oxford for helpful discussions about this project.

