## Supplementary Material for "Increased frequency of travel in the presence of cross-immunity may act to decrease the chance of a global pandemic"

**AUTHORS**

R.N. Thompson^1,2,3,^*, C.P. Thompson^2^, O. Pelerman^4^, S. Gupta^2^, U. Obolski^2,^*

**AFFILIATIONS**

^1^Mathematical Institute, University of Oxford, Andrew Wiles Building, Radcliffe Observatory Quarter, Woodstock Road, Oxford OX2 6GG, UK

^2^Department of Zoology, University of Oxford, South Parks Road, Oxford OX1 3PS, UK

^3^Christ Church, University of Oxford, St Aldate’s, Oxford OX1 1DP, UK

^4^The Chaim Rosenberg School of Jewish Studies, Tel Aviv University, Tel Aviv 69978, Israel

In our manuscript, we consider an outbreak of a high virulence (HV) pathogen strain which has the potential to cause a devastating pandemic. The HV strain outbreak has been preceded by an outbreak of a related low virulence (LV) strain of the pathogen. The population in which the outbreaks occur consists of two spatially-distinct subpopulations. Individuals infected in the LV strain outbreak are assumed to be partially immune against infection by the HV strain, as described in the main text. In the Supplementary Material, first we describe the model used to represent the LV strain outbreak, and consider the possible outcomes of that outbreak (Section 1). Then we consider the dynamics and possible outcomes of the HV strain outbreak (Section 2). These analyses include derivations of the probability of a major epidemic developing in each subpopulation, as well as the possible final sizes of the outbreaks in each subpopulation. We also present a table of the parameter values used in our analyses (Section 3), and compare our results which are based on quantities derived analytically or numerically with results generated using model simulations (Section 4). Finally, to supplement the Discussion in the main text, we conduct a crude estimation of travel rates from Europe to North America now compared with a century ago (Section 5), and perform a preliminary analysis of a scenario in which the HV strain outbreak can begin before the LV strain outbreak has ended (Section 6).

1. LV Strain Outbreak

Model

We consider a metapopulation model describing an outbreak of the LV strain in two spatially-distinct subpopulations each containing *N* individuals (Fig 1 of main text), with outbreak dynamics in each subpopulation governed by the stochastic Susceptible-Infected-Recovered (SIR) model. The deterministic SIR model in a single subpopulation consisting of *N* hosts is given by

$$\frac{dS}{dt}=-\beta_{LV}\kappa IS,$$

$$\frac{dI}{dt}=\beta_{LV}\kappa IS-\mu I,$$

$$\frac{dR}{dt}=\mu I. (S1)$$

The parameter $\beta_{LV}$ is the baseline infection rate of the LV strain. The parameter $\kappa$ is introduced as a proxy for the amount of within-subpopulation travel, relative to the baseline infection rate. For example, analyses in which $\kappa=2$ involve consideration of a population in which twice as many contacts occur per infected host every day compared to the baseline. Individuals are also assumed to travel between the subpopulations at a rate of $\lambda$ per day. In some of our analyses, we consider the analogous stochastic model, and when we run model simulations (Fig 3a, right column of Fig S1) we use the Gillespie direct method (Gillespie, 1977). We also approximate the behaviour of the stochastic model by deriving the probability of a major epidemic analytically, as well as approximating the expected final size.

Probability of a major epidemic (single subpopulation)

When the pathogen arrives in a subpopulation, the probability that the pathogen takes off and goes on to cause a major epidemic in that subpopulation can be derived by assuming that infections occur according to a branching process (see e.g. Keeling and Rohani, 2007). For the stochastic SIR model in a single subpopulation, this approximation to the probability of a major epidemic is given by

$$\mathbb{P}\left( major epidemic \right)\approx1-\left( \frac{1}{R_{0}^{LV}} \right)^{I(0)},$$

whenever $R_{0}^{LV}>1$ (otherwise the probability of a major epidemic is zero), where *I*(0) is the initial number of infected individuals and $R_{0}^{LV}=\frac{\beta_{LV}\kappa N}{\mu}$ is the within-subpopulation basic reproduction number of the pathogen.

Final size (single subpopulation)

If the pathogen takes off in a subpopulation and goes on to cause a major epidemic (which occurs according to the probability above), then dynamics are assumed to evolve according to the deterministic SIR model (equations (S1)) within that subpopulation. This allows us to calculate the expected final size, conditioned on a major epidemic occurring in that subpopulation.

To do this, we consider the system of equations (S1). Dividing the second equation by the first gives

$$\frac{dI}{dS}=-1+\frac{\mu}{\beta_{LV}\kappa S} ,$$

which has solution

$$I=-S+\frac{\mu}{\beta_{LV}\kappa}\ln S+C,$$

where *C* is a constant that can be found using the initial conditions, giving

$$I=-S+\frac{\mu}{\beta_{LV}\kappa}\ln S+S(0)+I(0)-\frac{\mu}{\beta_{LV}\kappa}\ln S(0).$$

Rearranging this expression, and noting that $S\left( 0 \right)+I\left( 0 \right)=N, I\left( \infty\right)=0$ and $R\left( \infty\right)=N-S(\infty)$, yields

$$R\left( \infty\right)=N-S\left( 0 \right)\exp\left( \frac{-R_{0}^{LV}R\left( \infty\right)}{N} \right). (S2)$$

The final size, $R\left( \infty\right)$ is therefore the solution of equation (S2).

Possible outcomes of LV strain outbreak (two subpopulations)

We now consider the possible outcomes of the LV strain outbreak in the full population consisting of two subpopulations. Since we are modelling a LV strain epidemic that confers partial cross-immunity against the HV strain when it arrives in the population, we assume that the LV strain outbreak is a major epidemic (i.e. there is a major epidemic of the LV strain in at least one of the two subpopulations).

The possible outcomes of the LV strain outbreak are therefore: i) a major epidemic in subpopulation 1 but not in subpopulation 2; ii) a major epidemic in subpopulation 2 but not in subpopulation 1; iii) a major epidemic in both subpopulations 1 and 2. Since the subpopulations that we consider are symmetric, we assume without loss of generality that the LV strain originated in subpopulation 1. We therefore neglect case ii, since the major epidemic in subpopulation 2 has to be seeded by individuals from subpopulation 1, thereby requiring a major epidemic in subpopulation 1 (we neglect the unlikely event that the small number of individuals infected in a minor outbreak in subpopulation 1 could travel to subpopulation 2 to cause a major epidemic in subpopulation 2).

Since a major epidemic occurs in subpopulation 1, by equation (S2) the final size of the LV strain outbreak in subpopulation 1 can be approximated by the solution $R^{LV}\left( \infty\right)$ of

$$R^{LV}\left( \infty\right)=N-N\exp\left( \frac{-R_{0}^{LV}R^{LV}\left( \infty\right)}{N} \right), (S3)$$

where we assume that the initial number of susceptible individuals in subpopulation 1 is $S\left( 0 \right)\approx N$.

If the LV strain has caused a major epidemic in subpopulation 1, then the probability of no major LV strain epidemic in subpopulation 2 is equal to the probability that *I** infected individuals transition to subpopulation 2 from subpopulation 1, and each fails to cause a major epidemic, summed over possible values of *I**. This is given by

$$\mathbb{P}\left( no major epidemic in subpopulation 2 \right)$$

$$=\sum_{i=0}^{R^{LV}\left( \infty\right)} \mathbb{P}\left( no major epidemic \right|I^{*}=i\mathbb{)P(}I^{*}=i),$$

$$\approx\sum_{i=0}^{R^{LV}\left( \infty\right)} \left( \frac{1}{R_{0}^{LV}} \right)^{i}{\binom{R^{LV}\left( \infty\right)}{i}\left( \frac{\lambda}{\lambda+\mu} \right)}^{i}\left( \frac{\mu}{\lambda+\mu} \right)^{R^{LV}\left( \infty\right)-i},$$

$$=\left( \frac{\lambda}{R_{0}^{LV}(\lambda+\mu)}+\frac{\mu}{\lambda+\mu} \right)^{R^{LV}\left( \infty\right)},$$

whenever $R_{0}^{LV}>1$. Assuming this is the case, the probability of a major epidemic of the LV strain in subpopulation 2, conditioned on a LV strain epidemic in subpopulation 1, is then given by

$$\mathbb{P}\left( major epidemic in subpopulation 2 \right)\approx1-\left( \frac{\lambda}{R_{0}^{LV}\left( \lambda+\mu\right)}+\frac{\mu}{\lambda+\mu} \right)^{R^{LV}\left( \infty\right)}. (S4)$$

If a major epidemic occurs in subpopulation 2, then the expected final size in that subpopulation depends on the number of infected individuals transitioning to subpopulation 2 from subpopulation 1, since this modifies the population size, *N*, in equation (S3). However, since the transition rate between subpopulations, $\lambda$, is small, we assume that only a small number of infected individuals transition from subpopulation 1 to subpopulation 2, so that *N* is approximately unchanged. Then, we can approximate the expected final size of the LV strain outbreak in subpopulation 2, conditioned on a major epidemic in subpopulation 2 occurring, using equation (S3).

In summary, we condition on a major epidemic of the LV strain occurring in subpopulation 1, with the final size in subpopulation 1 given by the solution of equation (S3). Then, the probability of a major epidemic of the LV strain in subpopulation 2 is given by equation (S4). If a major epidemic occurs in subpopulation 2, then the expected final size in that subpopulation is also given by the solution to equation (S3).

2. HV strain outbreak

Model

The outbreak of the HV strain follows the epidemic of the LV strain. Individuals infected in the LV strain epidemic are assumed to be partially immune against infection with the HV strain. This is implemented in the model of the HV strain outbreak by differentiating between individuals who were (*S*_I_) and were not (*S*_N_) previously infected by the LV strain. The initial fraction of susceptible individuals who were previously infected by the LV strain, which determines the initial value of $\frac{S_{I}}{S_{N}+S_{I}}$, is denoted by *x*.

Cross-immunity reduces the probability of infection compared to susceptible individuals who are not partially immune. The level of cross-immunity is governed by the parameter $\alpha$. The resulting deterministic S_N_S_I_IR model in a single subpopulation is then given by

$$\frac{dS_{N}}{dt}=-\beta_{HV}\kappa IS_{N},$$

$$\frac{dS_{I}}{dt}=-\beta_{HV}\kappa(1-\alpha)IS_{I},$$

$$\frac{dI}{dt}={\beta_{HV}\kappa IS_{N}+\beta}_{HV}\kappa\left( 1-\alpha\right)IS_{I}-\mu I,$$

$$\frac{dR}{dt}=\mu I. (S5)$$

Probability of a major epidemic (single subpopulation)

When the pathogen arrives in a subpopulation, the probability that the pathogen takes off and goes on to cause a major epidemic in that subpopulation can again be derived by assuming that infections occur according to a branching process. For the model in a single subpopulation governed by equations (S5), this approximation to the probability of a major epidemic is given by

$$\mathbb{P}\left( major epidemic \right)\approx1-\left( \frac{1}{R_{e}^{HV}} \right)^{I(0)},$$

whenever $R_{e}^{HV}>1$ (otherwise the probability of a major epidemic is zero), where *I*(0) is the initial number of infected individuals and $R_{e}^{HV}=\frac{\beta_{HV}\kappa(1-\alpha x)N}{\mu}$ is the effective reproduction number of the pathogen given that a proportion *x* of individuals in the subpopulation were previously infected by the LV strain. In our analyses, we set *I*(0) = 1. In the system that we consider, if a major epidemic of the LV strain previously occurred in the subpopulation concerned (so that *x* = $\frac{R^{LV}\left( \infty\right)}{N}$), then

$$R_{e}^{HV}=R_{e}=\frac{\beta_{HV}\kappa\left( 1-\alpha\frac{R^{LV}\left( \infty\right)}{N} \right)N}{\mu},$$

whereas if a major epidemic of the LV strain did not occur in the subpopulation concerned (so that $x\approx0$) then

$$R_{e}^{HV}=R_{0}^{HV}=\frac{\beta_{HV}\kappa N}{\mu}.$$

Final size (single subpopulation)

If the HV strain takes off and causes a major epidemic in a subpopulation, then it is assumed to follow deterministic dynamics within that subpopulation (governed by equations (S5)). This allows us to approximate the expected final size, conditioned on a major epidemic occurring in that subpopulation.

If a major epidemic of the LV strain has not previously occurred in the subpopulation, then since *S*_I_ = 0, equations (S5) reduce to the SIR model. The final size can then be approximated by the solution $R_{0}^{HV}$ of the final size equation

$$R_{0}^{HV}\left( \infty\right)=N-N\exp\left( \frac{-R_{0}^{HV}R_{0}^{HV}\left( \infty\right)}{N} \right).$$

If, on the other hand, a major epidemic of the LV strain has previously occurred in the subpopulation, then we solve equations (S5) numerically, starting with $S_{N}=(N-1)\left( 1-\frac{R^{LV}\left( \infty\right)}{N} \right)$, $S_{I}=(N-1)\frac{R^{LV}\left( \infty\right)}{N}$ , *I* = 1 and *R* = 0, to generate the expected number of individuals infected by the HV strain in that subpopulation, which we denote by $R_{e}^{HV}\left( \infty\right)$.

Possible outcomes of HV strain outbreak

There are a number of possible outcomes of the HV strain epidemic, depending on the prior outcome of the LV strain epidemic. The overall possible outcomes of the LV and HV strain outbreaks, conditioned on a major epidemic of the LV strain in subpopulation 1, are given by

| **Case** | **Major epidemic of LV strain in subpopulation 1** | **Major epidemic of LV strain in subpopulation 2** | **Major epidemic of HV strain in subpopulation 1** | **Major epidemic of HV strain in subpopulation 2** |
| --- | --- | --- | --- | --- |
| 1 | YES | YES | NO | NO |
| 2 | YES | YES | YES | NO |
| 3 | YES | YES | NO | YES |
| 4 | YES | YES | YES | YES |
| 5 | YES | NO | NO | NO |
| 6 | YES | NO | YES | NO |
| 7 | YES | NO | NO | YES |
| 8 | YES | NO | YES | YES |

**Table S1.** The possible outcomes of the outbreaks of the LV and HV strains. We assume that the LV strain outbreak arrives in subpopulation 1 and that a major epidemic of the LV strain follows.

We assume that the HV strain outbreak is equally likely to be seeded in each subpopulation. The associated probabilities of each of the cases above occurring can be approximated analytically.

For example, for case 1 to occur, the LV epidemic in subpopulation 1 has to drive a major epidemic in subpopulation 2 (see equation (S4) for the associated probability), and the HV epidemic has to fail to take off when the HV strain arrives into the population. The resulting probability is then

$$\mathbb{P}\left( Case 1 \right)=\left( 1-\left( \frac{\lambda}{R_{0}^{LV}(\lambda+\mu)}+\frac{\mu}{\lambda+\mu} \right)^{R^{LV}\left( \infty\right)} \right)\cdot\left( \frac{1}{R_{e}} \right).$$

Case 2 can only occur if the HV strain arrives in subpopulation 1. Then, a major epidemic must be seeded (with probability 0.5) and arise in subpopulation 1, and no major epidemic must occur in subpopulation 2, so that

$$\mathbb{P}\left( Case 2 \right)=\left( 1-\left( \frac{\lambda}{R_{0}^{LV}(\lambda+\mu)}+\frac{\mu}{\lambda+\mu} \right)^{R^{LV}\left( \infty\right)} \right)\times\frac{1}{2}\times\left( 1-\frac{1}{R_{e}} \right)\times\left( \left( \frac{\lambda}{R_{e}(\lambda+\mu)}+\frac{\mu}{\lambda+\mu} \right)^{R_{e}^{HV}\left( \infty\right)} \right),$$

assuming that ${R_{0}^{LV}, R}_{e}>1$ (otherwise the probability of case 2 occurring is zero). The probabilities associated with each of the other cases can be calculated similarly.

The probability of a major epidemic of the HV strain (defined as a major epidemic of the HV strain in either subpopulation, or in both subpopulations) can then be found by summing the relevant probabilities,

$\mathbb{P}\left( major epidemic of HV strain \right)=\mathbb{P}\left( Case 2 \right)\mathbb{+P}\left( Case 3 \right)\mathbb{+P}\left( Case 4 \right)\mathbb{+P}\left( Case 6 \right)\mathbb{+ P}\left( Case 7 \right)\mathbb{+P}\left( Case 8 \right)$.

After some simplification, this reduces to

$$\mathbb{P}\left( major epidemic of HV strain \right)=1-\frac{1}{R_{e}}+\frac{1}{2}\left( \frac{\lambda}{R_{0}^{LV}\left( \lambda+\mu\right)}+\frac{\mu}{\lambda+\mu} \right)^{R^{LV}\left( \infty\right)}\left( \frac{1}{R_{e}}-\frac{1}{R_{0}^{HV}} \right),$$

whenever ${R_{0}^{LV}, R}_{e}>1$.

The expected final size of the HV strain epidemic can also be calculated,

$$\mathbb{E}\left( final size of HV strain epidemic \right)=\sum_{j=1}^{8} \mathbb{E}\left( final size of HV strain epidemic | Case j \right)\mathbb{P}\left( \mathrm{Case}j \right),$$

as well as the expected final size of the HV strain epidemic conditioned on a major epidemic of the HV strain,

$$\mathbb{E}\left( final size of HV strain epidemic | major epidemic of HV strain \right)=\frac{\sum_{j=2,3,4,6,7,8} \mathbb{E}\left( final size of HV strain epidemic | Case j \right)\mathbb{P}\left( \mathrm{Case}j \right)}{\sum_{j=2,3,4,6,7,8} \mathbb{P}\left( \mathrm{Case}j \right)}.$$

3. Table of parameters

We present a table of the baseline parameter values used in our analyses below. These values are used except where stated in the figure captions.

| **Parameter** | **Definition** | **Default value (except where stated)** |
| --- | --- | --- |
| *N* | Effective population size in each subpopulation | 1000 |
| $\kappa$ | Extent of within-subpopulation travel | 1 |
| $\beta_{LV}$ | Infection rate parameter for LV strain | 2.86 x 10^-4^ day^-1^ (chosen so that$R_{0}^{LV}=2$ when $\kappa=1$) |
| $\beta_{HV}$ | Infection rate parameter for HV strain | 4.29 x 10^-4^ day^-1^ (chosen so that$R_{0}^{HV}=3 \mathrm{when}\kappa=1$) |
| $\mu$ | Recovery/death rate | 0.14 day^-1^ |
| $\lambda$ | Migration rate between subpopulations | See figures |

*The following quantities are then derived:*

| $R_{0}^{LV}$ | Within-subpopulation basic reproduction number of LV strain | 2 ($\mathrm{when}\kappa=1$) |
| --- | --- | --- |
| $R_{0}^{HV}$ | Within-subpopulation basic reproduction number of HV strain | 3 ($\mathrm{when}\kappa=1$) |
| $R_{e}$ | Within-subpopulation effective reproduction number of HV strain, given cross-immunity | 1.33 ($\mathrm{when}\kappa=1$) |
| $R^{LV}(\infty)$ | Final size in a single subpopulation of a major epidemic of the LV strain | 797 ($\mathrm{when}\kappa=1$) |
| $R_{0}^{HV}(\infty)$ | Final size of a major epidemic of the HV strain in a subpopulation in which a major epidemic of the LV strain has not occurred | 940 ($\mathrm{when}\kappa=1$) |
| $R_{e}^{HV}(\infty)$ | Final size of a major epidemic of the HV strain in a subpopulation in which a major epidemic of the LV strain has occurred | 342 ($\mathrm{when}\kappa=1$) |

**Table S2.** The parameters governing disease transmission in the two-subpopulation model, along with their default values.

4. Verifying the analytic results

In our analyses with results displayed in the main text (except Fig 3), we derived estimates of quantities pertaining to the outbreaks of the LV and HV strains, such as the probability of a major epidemic occurring in each subpopulation, and the expected final sizes conditioned on a major epidemic. However, we also checked that the results in the main text matched the mean results of stochastic simulations generated using the Gillespie direct method (Gillespie, 1977). As an example, in Fig S1 we display results from simulations corresponding to the middle column of Fig 4 from the main text. For computational efficiency due to the large number of simulations required, in Fig S1 we only discretise the contour plot into boxes of width (change in $\kappa$) 0.1 and height (change in $\lambda$) 5 x 10^-5^ day^-1^.

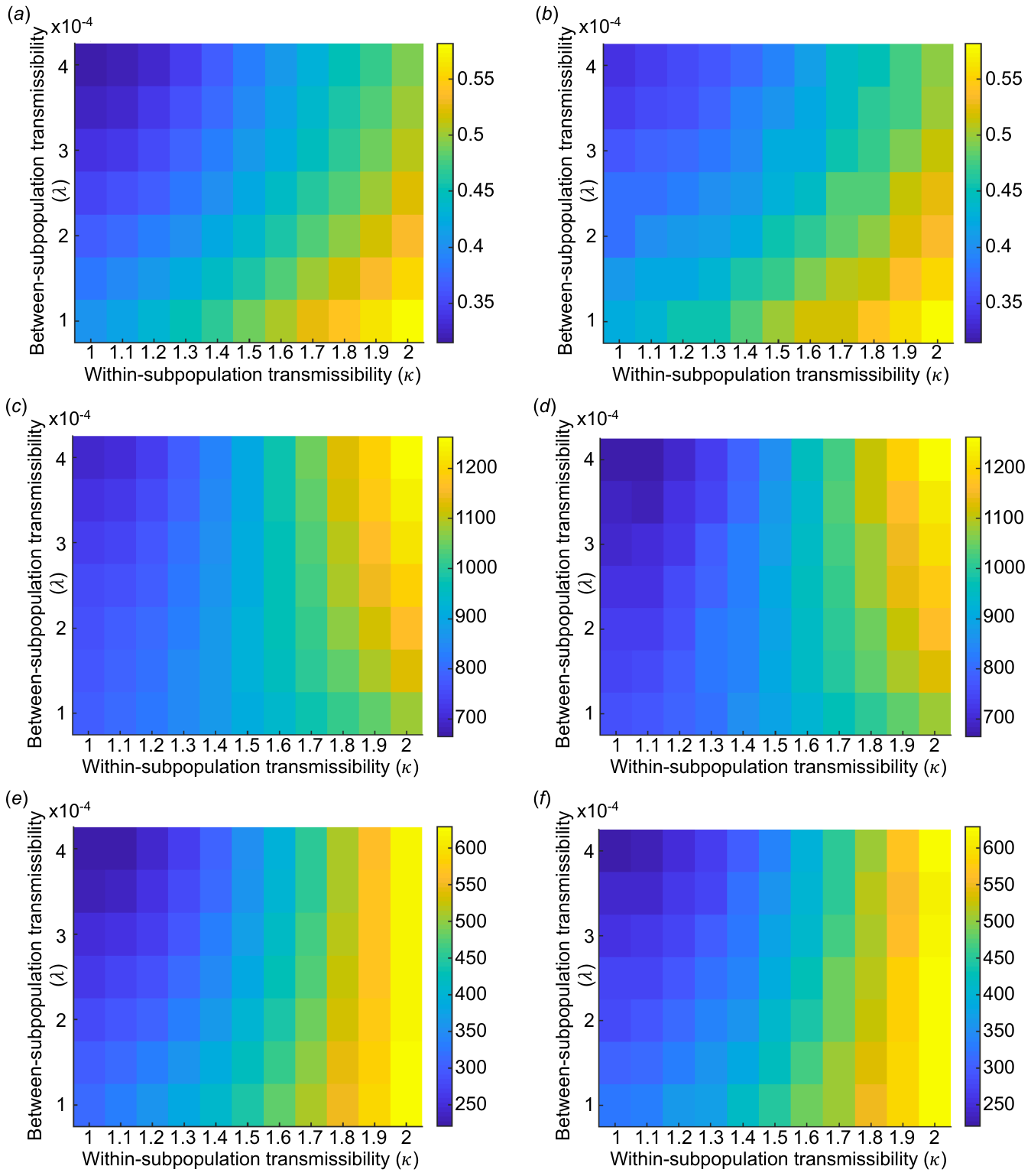

**Figure S1.** Verifying that our analyses match the results of stochastic simulations. Left column shows our analytic results (equivalent to middle panel of Fig 4 in the main text, but with fewer boxes). Right column shows the analogous results obtained using stochastic simulations, with 1000 simulations per box. (a)-(b): the probability of a major epidemic of the HV strain. (c)-(d): the final size of a major epidemic of the HV strain. (e)-(f): the final size of an outbreak of the HV strain, allowing for the possibility of a major epidemic or fade out without causing a major epidemic. Parameter values: $N=1000$ individuals per subpopulation,$\alpha=0.7$,$R_{0}^{LV}=2\kappa,R_{0}^{HV}=3\kappa$. For a description of the parameters, see Table S2.

5. Travel rates estimates

In the Discussion of our manuscript, we approximated the daily travel rates of individuals from Europe to the USA in the early 20^th^ century and in the present day. To do this, we obtained data on the population size, *N*, and the annual number of transatlantic journeys, *Y*, in each of the years in the period 1914-1924 as well as in 2016. The daily per capita travel rates were then calculated as follows.

First, the probability of an individual travelling from Europe to USA in each year was approximated as $\frac{Y}{N}$. Then, the daily probability is $\frac{Y}{365\cdot N}$. Assuming that the periods between travel events, which occur at rate $\lambda$ per day, are exponentially distributed, then the probability of travel per day is given by

$$1-e^{-\lambda}=\frac{Y}{365\cdot N},$$

which can be rearranged to give

$$\lambda=-log\left( 1-\frac{Y}{365\cdot N} \right).$$

The data underlying this calculation (Data S1) were obtained from a number of sources. The population sizes of countries (corresponding approximately to the modern-day European Union) from 1914-1924 were obtained from Lahmeyer (2018). The numbers of transatlantic journeys in each of these years were found in port registry data (Wilcox *et al*., 1931). For the present-day calculation, we used data on the total population size living in the European Union in 2016 (Eurostat, 2018a), as well as the annual number of departing passengers from the European Union to the USA (Eurostat, 2018b). Our calculations can be found in Data S1.

6. Timing of introduction of HV strain

In the analyses in the main text, we assumed that the LV strain epidemic had ended when the HV strain appeared in the system. This is characteristic of, for example, influenza epidemics in different seasons. However, for some pathogens, the HV strain might enter the system while the LV strain is still circulating. We therefore conducted a supplementary analysis, in which the timing of the introduction of the HV strain was varied.

We extended the epidemiological model to allow the HV strain to circulate before the end of the LV strain epidemic. We assumed that individuals could only be infected by a single strain at any time, and that individuals infected by the HV strain could not subsequently be infected by the LV strain. We also assumed that individuals infected with the LV strain could be infected by the HV strain, but that in that case the HV strain infection immediately took over from the less virulent LV strain infection, thereby maintaining each individual only infected by at most a single strain at any time. We assumed that the LV strain provided complete protection against reinfection with the same strain. The deterministic model that is analogous to the stochastic model we then considered is given by

$$\frac{dS_{N}}{dt}=-\beta_{HV}\kappa I_{HV}S_{N}-\beta_{LV}\kappa I_{LV}S_{N},$$

$$\frac{dS_{I}}{dt}=-\beta_{HV}\kappa\left( 1-\alpha\right)I_{HV}S_{I}+\mu I_{LV},$$

$$\frac{dI_{LV}}{dt}=\beta_{LV}{\kappa I}_{LV}S_{N}-\beta_{HV}\kappa\left( 1-\alpha\right)I_{HV}I_{LV}-\mu I_{LV},$$

$$\frac{dI_{HV}}{dt}={\beta_{HV}\kappa I_{HV}S_{N}+\beta}_{HV}\kappa\left( 1-\alpha\right)I_{HV}S_{I}+\beta_{HV}\kappa\left( 1-\alpha\right)I_{HV}I_{LV}-\mu I_{HV},$$

$$\frac{dR}{dt}=\mu I_{HV}.$$

In this model, individuals in the $S_{N}$ class have not been infected by either strain and so are fully susceptible to both strains, whereas individuals in the $S_{I}$ class have previously been infected by the LV strain, and so are fully immune against reinfection by the LV strain and partially cross-immune against infection by the HV strain. The infection rate of individuals in the $S_{I}$ class by the HV strain is identical to the infection rate of individuals in the $I_{LV}$ class by the HV strain.

We ran stochastic simulations of the LV strain epidemic, and introduced a single individual infected with the HV strain after *T* days. We ran 100,000 simulations for each possible value of *T* = 0,1,2,…,150 days, and recorded the probability of a major epidemic of the HV strain (Fig S2a) – defined as outbreaks in which there are more than 20 cases of HV strain infections – and the mean final size of all HV strain outbreaks (Fig S2b). We found that, the later the HV strain arrived in the system, the lower the chance of a major epidemic and the smaller the final size. This is because, if the HV strain arrives later, the LV strain has had more opportunity to spread through the population, thereby leading to a larger number of individuals likely to be cross-immune against the HV strain.

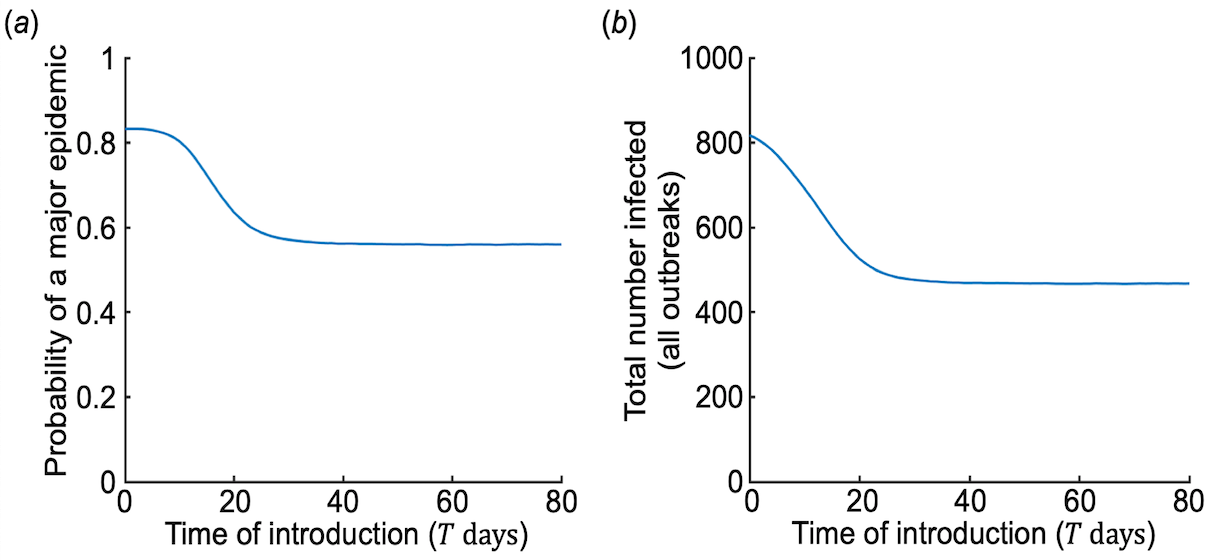

**Figure S2.** The probability of a major epidemic of the HV strain, and the size of the HV strain outbreak, when the HV strain can arrive in the population before the end of the LV strain outbreak. (a) The probability of a major epidemic of the HV strain, for different times of introduction after the start of the LV strain outbreak (*T* days). (b) The mean size of the HV strain epidemic, for different times of introduction after the start of the LV strain outbreak (*T* days). In this analysis, so that the results are comparable for different values of *T*, we do not condition of the LV strain outbreak being a major epidemic (which is affected by the value of *T*). Parameter values: $N=1000$ individuals per subpopulation,$\alpha=0.7$,${\kappa=2, R}_{0}^{LV}=2\kappa,R_{0}^{HV}=3\kappa$. For a description of the parameters, see Table S2.
